# A SUMO-Ubiquitin Relay Recruits Proteasomes to Chromosome Axes to Regulate Meiotic Recombination

**DOI:** 10.1101/095711

**Authors:** H.B.D. Prasada Rao, Huanyu Qiao, Shubhang K. Bhatt, Logan R.J. Bailey, Hung D. Tran, Sarah L. Bourne, Wendy Qiu, Anusha Deshpande, Ajay N. Sharma, Connor J. Beebout, Roberto J. Pezza, Neil Hunter

## Abstract

Meiosis produces haploid gametes through a succession of chromosomal events including pairing, synapsis and recombination. Mechanisms that orchestrate these events remain poorly understood. We found that the SUMO-modification and ubiquitin-proteasomes systems regulate the major events of meiotic prophase in mouse. Interdependent localization of SUMO, ubiquitin and proteasomes along chromosome axes was mediated largely by RNF212 and HEI10, two E3 ligases that are also essential for crossover recombination. RNF212-dependent SUMO conjugation effected a checkpoint-like process that stalls recombination by rendering the turnover of a subset of recombination factors dependent on HEI10-mediated ubiquitylation. We propose that SUMO conjugation establishes a precondition for designating crossover sites via selective protein stabilization. Thus, meiotic chromosome axes are hubs for regulated proteolysis via SUMO-dependent control of the ubiquitin-proteasome system.

**One Sentence Summary:** Chromosomal events of meiotic prophase in mouse are regulated by proteasome-dependent protein degradation.

---

Meiosis halves the chromosome complement via two successive rounds of cell division. Accurate segregation of homologous chromosomes (homologs) during the first meiotic division requires their connection by chiasmata — the conjunction of crossing over and sister-chromatid cohesion (*1*). Chiasma formation is the culmination of an elaborate series of interdependent events that include programmed recombination, and the pairing and synapsis of homologs. Each homolog comprises two sister chromatids, organized into arrays of chromatin loops connected to a common core or axis. Pairing and synapsis are promoted by homologous recombination, which occurs in physical and functional association with these axes. As meiosis progresses, axes align and become connected along their lengths to form synaptonemal complexes (SCs).

The SUMO-modification (SMS) and ubiquitin-proteasome (UPS) systems are key regulators of cellular proteostasis (*2*, *3*) and are implicated in various aspects of meiotic prophase (*4*–*9*). However, their roles remain poorly characterized, especially in mammalian meiosis. To obtain cytological evidence that the SMS and UPS regulate axis-associated events, we analyzed the localization of SUMO, ubiquitin and proteasomes along surface-spread chromosomes from mouse spermatocytes (Fig. 1 and figs. S1-S3).

**Fig. 1.**
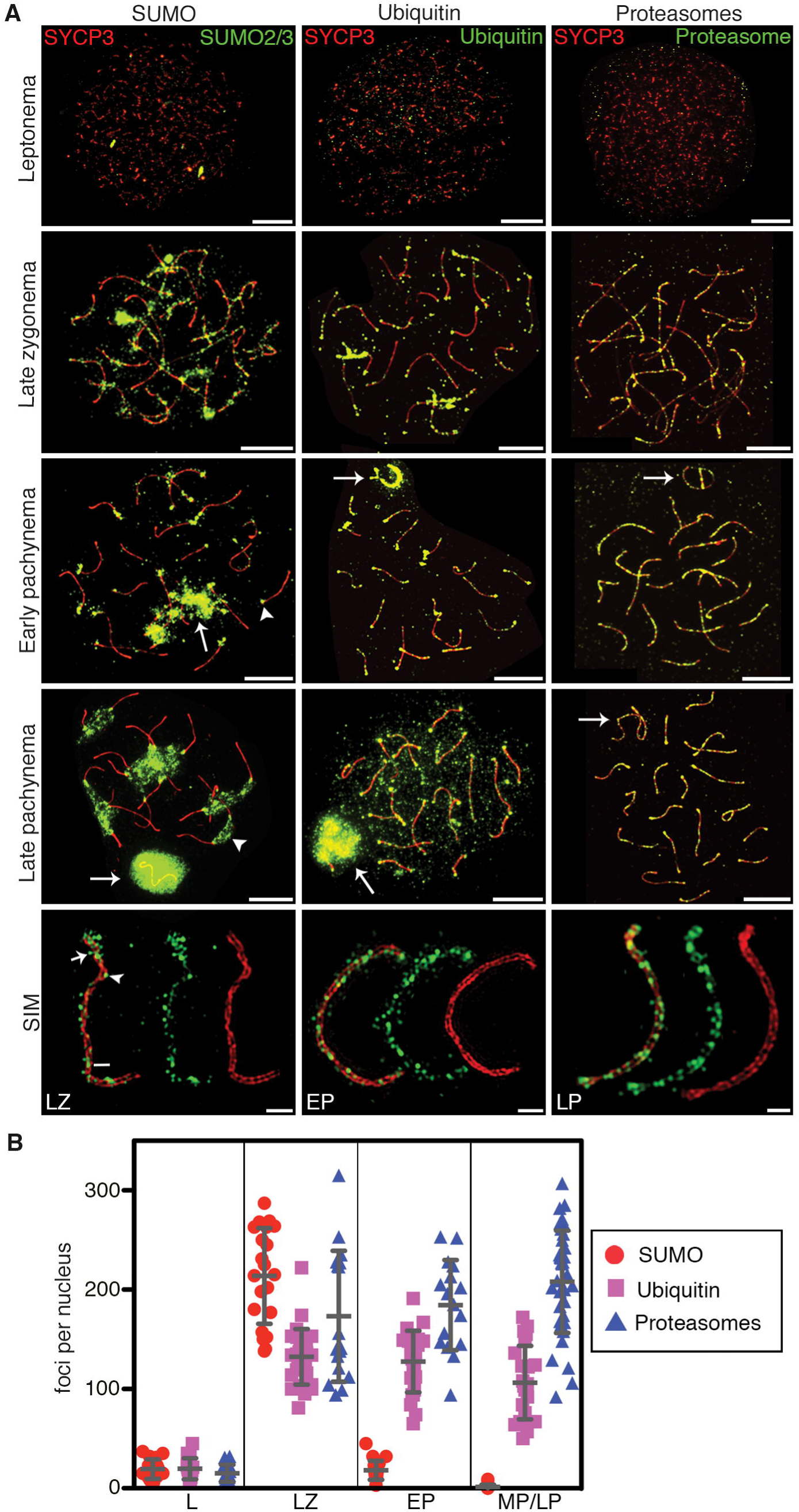
SUMO, ubiquitin and proteasomes decorate the axes of meiotic prophase chromosomes. (**A**) Spermatocyte nuclei immunostained for axis marker SYCP3 and either SUMO2/3, ubiquitin or proteasomes. Bottom row shows single chromosomes imaged by SIM. Arrows indicate sex bodies; arrowheads highlight centromeric heterochromatin. In the SUMO2/3 stained SIM image, the arrow, arrowhead and bar highlight supra-axial, axial and SC central region localizations, respectively. Scale bars: 10μm for widefield, 1μm for SIM. (**B**) Quantification of immunostaining foci. Leptonema (L), late zygonema (LZ), early pachynema (EP), and mid/late pachynema (MP/LP). Counts from MP and LP nuclei were pooled as numbers remained stable over these stages. Ubiquitin foci in LP could not be accurately quantified because general chromatin staining obscured axis-specific foci. Error bars show mean ± s.d.

The SUMO1 and SUMO2/3 isoforms localized to axes during zygonema, as chromosomes underwent synapsis, forming punctate patterns of ~200 immunostaining foci (Fig. 1A,B and fig. S1). Super-resolution Structured Illumination Microscopy (SIM) revealed that SUMO was present on both unsynapsed and synapsed axes; axial, supra-axial (extending into adjacent chromatin) and SC central-region staining could be discerned (Fig. 1A and fig. S1). General axis staining disappeared after synapsis completed and cells entered pachynema (Fig. 1A,B and fig. S1). Subsequently, SUMO accumulated on centromeric heterochromatin and the XY chromatin (sex body) (*10*). Prominent axis staining of ubiquitin was also detected during zygonema, but persisted throughout pachynema (Fig. 1A,B and fig. S2). By SIM, most ubiquitin foci localized to axes, but SC central region staining was occasionally discerned (Fig. 1A and fig. S2). Ubiquitin also accumulated along axes of the sex chromosomes during zygonema and early pachynema, before spreading to the entire XY chromatin (*11*). Prominent staining of centromeric heterochromatin was not seen for ubiquitin, but general chromatin staining became apparent after mid pachynema (Fig. 1A,B and fig. S2). Abundant recruitment of proteasomes along axes also occurred during zygonema (Fig. 1A,B and fig. S3), and persisted throughout pachynema and diplonema, when chromosomes desynapsed. By SIM, proteasome foci were largely axis associated, but less frequent SC central-region staining was also seen. Sex body, centromeric and general chromatin staining were not detected. This sub-chromosomal recruitment of proteasomes to meiotic chromosome axes, which appears to be an evolutionarily conserved feature of meiosis (see accompany paper from Ahuja et al.), predicts that axis-associated ubiquitin should include K48-linked chains; this inference was confirmed using linkage-specific ubiquitin antibodies (fig. S2)(*12*).

These immunostaining results suggest that the SMS and UPS regulate axis-associated events via protein degradation. To test this, we exploited short-term culture of testis cell suspensions (*13*) and chemical inhibitors of SUMO conjugation (2-DO8)(*14*), ubiquitin activation (PYR41)(*15*) or proteasomal degradation (MG132) (Fig. 2 and fig. S4-S6). All three inhibitors caused dramatic increases in large extra-chromosomal aggregates containing SYCP3 and SYCP2, two meiosis-specific components of chromosome axes (*16*)(Fig. 2A,B and fig. S4). Large increases in synapsis defects were also observed (Fig. 2A,C and fig. S4). Thus, the SMS and UPS modulate axis assembly and formation/maintenance of synapsis in mammalian meiosis. Phenotypes of *S.cerevisiae* and *C. elegans* mutants defective for proteasome (see accompany paper from Ahuja et al.), or CSN/COP9 signalosome (*17*) functions imply that a role for the UPS in synapsis is likely to be conserved.

**Fig. 2.**
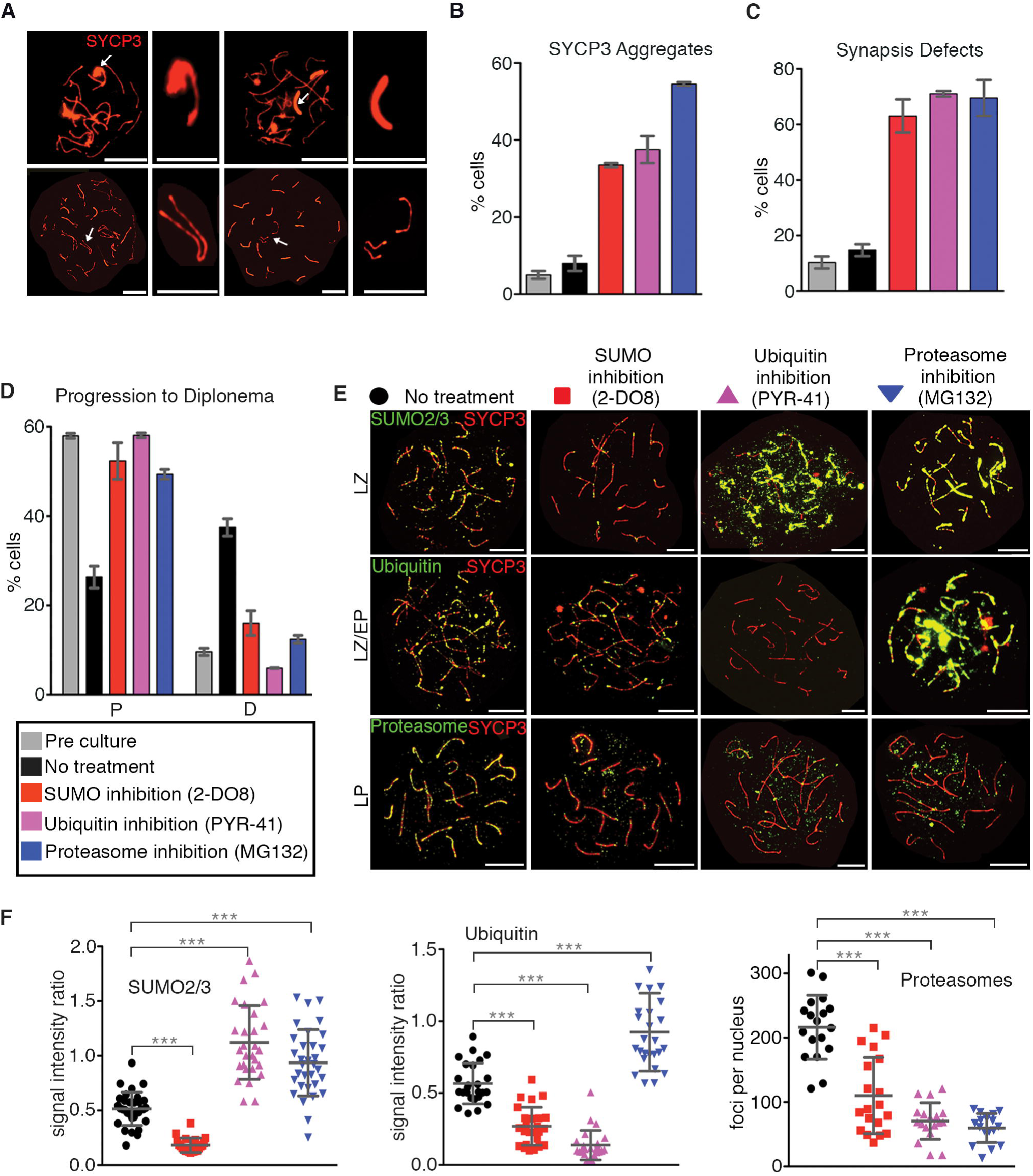
Interrelated functions of the SMS and UPS in meiotic prophase. (**A**) Pachytene-like spermatocyte nuclei containing SYCP3 aggregates (upper panels) or synapsis defects (lower panels). Arrows indicate structures magnified in adjacent panels. (**B** and **C**) Quantification of SYCP3 aggregates (**B**) and synapsis defects (**C**). (**D**) Progression of spermatocytes following culture with or without inhibitors. (**E**) Spermatocyte nuclei immunostained for SYCP3 (red), and either SUMO2/3, ubiquitin, or proteasomes (green) following chemical inhibition. (**F**) Quantification of experiments shown in (E). Error bars in B, C and D show mean ± s.e.m. Error bars in G show mean ± s.d. (***) *P* ≤ 0.001, two-tailed Mann-Whitney. LZ, late zygonema; MP, mid pachynema; LP, late pachynema. Scale bars: 10 μm for full nuclei and 1μm for magnified structures in A.

Absent inhibitors, cultured spermatocytes continued to progress through meiotic prophase (*13*), manifested by a reduction in pachytene cells and a corresponding increase in diplotene cells after 24 hours (Fig. 2D). Treatment with 2-DO8, PYR41 or MG132 blocked progression out of pachynema (Fig. 2D). Moreover, precocious progression into metaphase-I triggered by phosphatase inhibitor, okadaic acid, was blocked by 2-DO8, PYR41 and MG132 (**fig. S5**). Thus, the SMS and UPS are important for both entry and exit from the pachytene stage of meiosis.

A dependent relationship between the SMS and UPS was revealed by immunostaining analysis of inhibitor-treated spermatocytes. 2-DO8 diminished not only SUMO staining (Fig. 2E,F and **fig. S6**), but also axis-associated ubiquitin and proteasomes (Fig. 2E,F), raising the possibility that SUMO-modified proteins become substrates for the UPS. Indeed, SUMO staining accumulated and persisted when ubiquitylation was inhibited, while proteasome recruitment remained defective (Fig. 2E,F and **fig. S6**). Furthermore, chromosomal staining of both SUMO and ubiquitin became abundant and persistent when proteasomes were inhibited (Fig. 2E,F and **fig. S6**).

Effects of inhibiting the SMS and UPS on recombination were investigated by immunostaining for markers representing each step of the pathway (Fig. 3 and **figs. S7, S8**). MEI4 is required for initiation of recombination by DNA double-strand breaks (DSBs) and localizes as several hundred foci along axes during leptonema and early zygonema (*18*)(Fig. 3A). In untreated spermatocytes, MEI4 foci decreased between early and mid zygonema (Fig. 3B). Inhibition of SUMO, ubiquitin or proteasomes did not alter MEI4 foci in early zygonema, but numbers remained aberrantly high in mid zygonema and, to a lesser extent, late zygonema (Fig. 3A,B) implying that the SMS and UPS promote MEI4 turnover. RAD51 is a RecA-family protein that promotes DNA pairing and strand exchange (*19*). In control cells, RAD51 foci were abundant during early zygonema and diminished throughout zygonema as recombination proceeded (Fig. 3C,D and **fig. S7**). Initial numbers of RAD51 foci were not changed by drug treatments suggesting that DSB numbers and assembly of RAD51 complexes were unaffected; however, turnover of RAD51 was defective. SUMO inhibition decreased RAD51 foci in mid-zygonema by ~61% implying accelerated turnover (Fig. 3C,D and **fig. S7**). In contrast, when ubiquitylation or proteasomes where inhibited, RAD51 foci remained high, consistent with defective turnover (Fig. 3C,D and **fig. S7**). Analogous results were obtained for meiosis-specific RecA homolog, DMC1 (**fig. S7**).

**Fig. 3.**
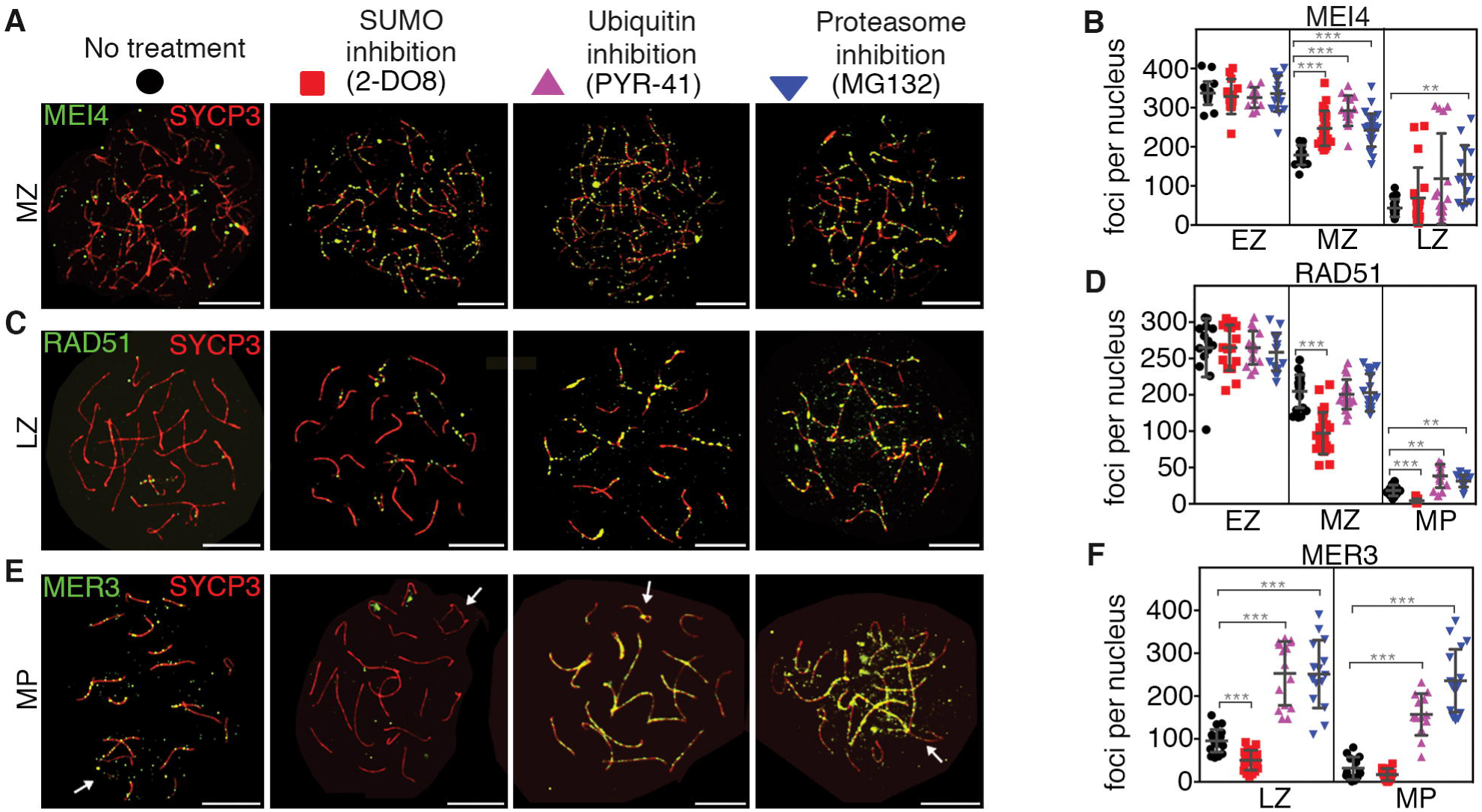
The SMS and UPS regulate chromosomal dynamics of recombination factors. (**A, C, E** and **G**) Spermatocyte nuclei immunostained for SYCP3 (red) and either MEI4, RAD51, or MER3 (green) following chemical inhibition. (**B, D, F** and **H**) Quantification of immunostaining foci. EZ, early zygonema; MZ, mid zygonema; LZ, late zygonema; MP, mid pachynema; LP, late pachynema. Arrows indicates sex chromosomes. Error bars show mean ± s.d. Scale bars = 10μm.

As chromosomes synapse, a distinct set of recombination factors associate with homolog axes, including ZMM proteins, which are required for efficient synapsis and crossing over (*1*). DNA helicase, MER3, stabilizes nascent DNA strand exchange intermediates. Inhibition of the SMS or UPS had dramatically different effects on MER3 dynamics: SUMO inhibition halved the numbers of MER3 foci in late zygotene nuclei, while ubiquitin and proteasome inhibition increased MER3 foci ~2.5-fold (Fig. 3E,F). Similar trends were seen for three other ZMM members (TEX11, MSH4 and RNF212) and ssDNA-binding protein RPA, which localizes to recombination sites with timing similar to the ZMMs (**fig. S8**). Finally, we analyzed the localization of ubiquitin-ligase, HEI10, which specifically marked crossover sites during mid-late pachynema (Fig. 3G,H)(*20*). HEI10 focus numbers increased only very slightly following inhibition of the SMS or UPS (Fig. 3G,H). However, SUMO inhibition resulted in significantly brighter HEI10 foci (**fig. S9**), implying that SUMOylation negatively regulates HEI10 accumulation. Together, inhibitor studies indicate general roles for the SMS and UPS in regulating turnover of recombination factors; and more specifically reveal opposing effects on the chromosomal dynamics of ZMMs and associated factors, with stabilization being promoted by the SMS and turnover by the UPS. This interpretation is supported by genetic analysis below.

Two RING-family E3 ligases are essential for crossing over in mouse (*20*, *21*). RNF212 is inferred to catalyze SUMO conjugation and promotes selective stabilization of ZMM factor MutSγ at the minority of recombination sites that will mature into crossovers (~10% in mouse)(*21*). Oppositely, HEI10 stimulates ubiquitylation and antagonizes RNF212 by promoting its turnover from synapsed chromosomes (*20*). To understand the relationships between RNF212, HEI10 and the axis-localized SUMO, ubiquitin and proteasomes described here, immunostaining was performed on spermatocyte chromosomes from *Rnf212^-/-^* and *Hei10^-/-^* single mutant mice, and the *Rnf212^-/-^ Hei10^-/-^* double mutant (Fig. 4; in subsequent analyses, spermatocyte chromosome spreads were prepared directly from testes, without intervening cell culture). Axis SUMOylation was largely dependent on RNF212 (Fig. 4A,B and **fig. S10**). However, accumulation of SUMO at the sex body and centromeric heterochromatin were still observed in *Rnf212^-/-^* nuclei (**fig. S10**), indicating that RNF212 promotes SUMO conjugation specifically along chromosome axes. Oppositely, SUMO was abundant and persistent along synapsed chromosomes in *Hei10^-/-^* spermatocytes, with high levels now being detected throughout pachynema (Fig. 4A,B and **fig. S10**). Analysis of the *Rnf212^-/-^ Hei10^-/-^* double mutant revealed that this hyper-SUMO phenotype was RNF212 dependent (Fig. 4A,B). Thus, RNF212 and HEI10 mediate, respectively, formation and turnover of axis-associated SUMO conjugates.

**Fig. 4.**
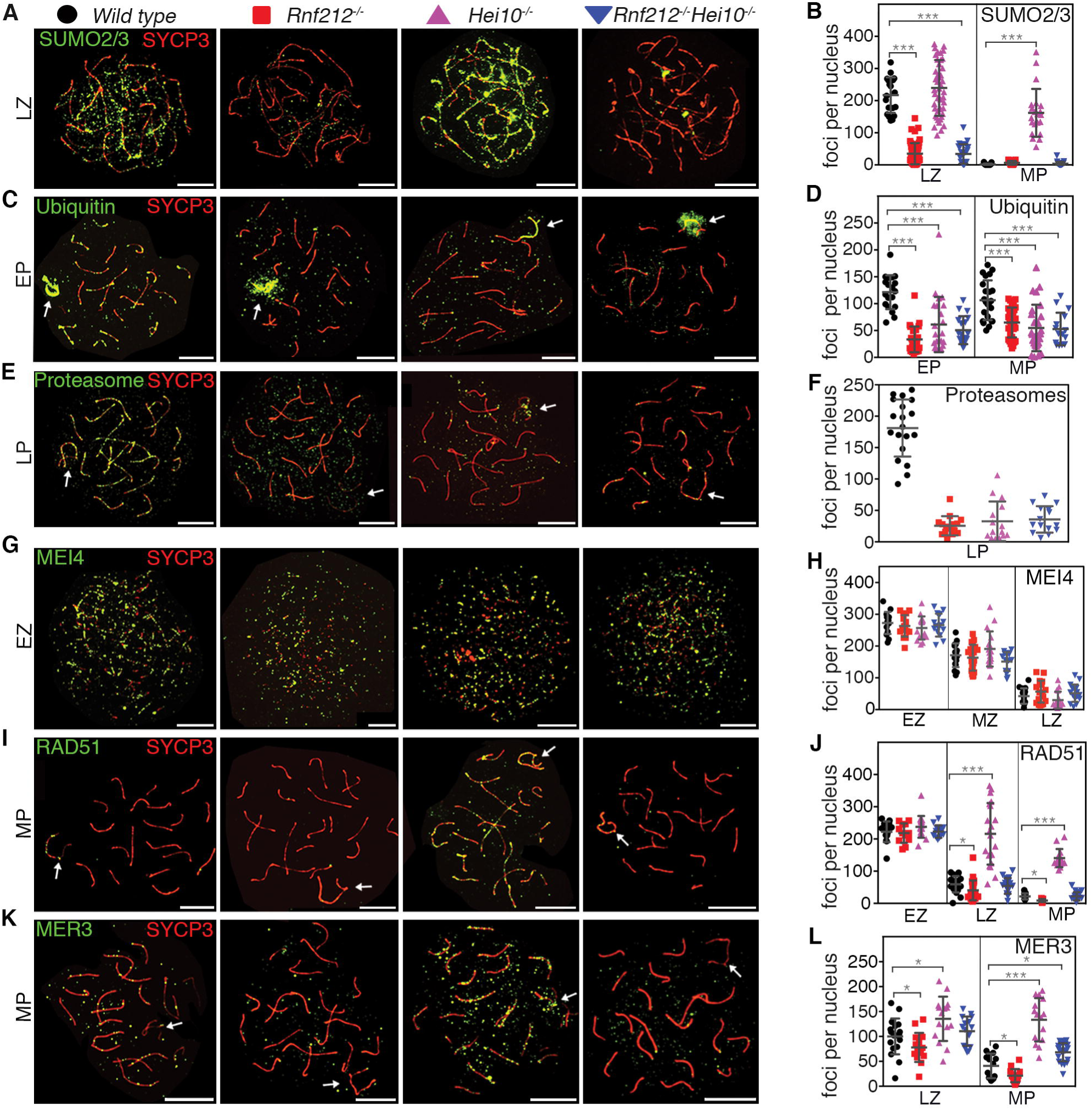
RNF212 and HEI10 define an axis-associated SUMO-ubiquitin-proteasome relay that regulates turnover of recombination factors. (**A,C,E**) Spermatocyte nuclei from indicated strains immunostained for SYCP3 (red) and either SUMO2/3, ubiquitin, or proteasomes (green). See fig. S8 for SUMO1 staining. (**B,D,F**) Quantification of immunsoatining foci. (**G,I,K**) Spermatocyte nuclei immunostained for SYCP3 (red) and either MEI4, RAD51 or MER3 (green). (**H,J,L**) Quantification of immunsoatining foci. EZ, early zygonema; LZ, late zygonema; MP, mid pachynema. Arrows indicate sex chromosomes. Error bars show mean ± s.d. (*) *P* ≤ 0.05, (***) *P* ≤ 0.001, two-tailed Mann-Whitney. Scale bars = 10μm.

Consistent with a model in which RNF212-mediated SUMO conjugation creates substrates for HEI10-dependent proteasomal degradation, axis-localized ubiquitin and the recruitment of proteasomes were diminished in *Rnf212^-/-^* and *Hei10^-/-^* single mutants, and the *Rnf212^-/-^ Hei10^-/-^* double mutant (Fig. 4C-F and **fig. S10**; ubiquitin staining of sex chromosomes appeared unaffected). Analogous observations were made for female meiosis (**fig. S11**). Together, these results point to the same pathway relationships inferred from inhibitor studies, with RNF212-dependent SUMOylation establishing a requirement for ubiquitin-dependent proteolysis via HEI10. However, unlike general inhibition of the SMS and UPS, *Rnf212^-/-^* and *Hei10^-/-^* mutations do not cause overt defects in synapsis or progression to metaphase (*20*, *21*). Thus, the SUMO-ubiquitin-proteasome relay defined by RNF212 and HEI10 appears dedicated to the regulation of post-synapsis steps of meiotic recombination.

To better understand how recombination is regulated by the RNF212-HEI10 pathway, we analyzed the same set of markers described above for chemical inhibition experiments (Fig. 4G-L and **figs. S12, S13**). While pan-inhibition caused abnormally high numbers of MEI4 foci to persist through zygotene (Fig. 3A,B), mutation of *Rnf212* and/or *Hei10* did not (Fig. 4G,H) indicating that MEI4 turnover does not require the RNF212-HEI10 pathway. In contrast, RAD51 turnover was strongly dependent on HEI10; in *Hei10^-/-^* nuclei, high numbers of RAD51 foci persisted throughout zygonema and pachynema (Fig. 4I,J and **fig. S12**). Moreover, persistence of RAD51 was RNF212 dependent as foci decreased to wild-type levels in the *Rnf212^-/-^ Hei10^-/-^* double mutant. Unlike global UPS inhibition, which impeded turnover of both RAD51 and DMC1, *Hei10^-/-^* mutation did not slow the disappearance of DMC1 foci (**fig. S12**). Thus, turnover of the two RecA homologs involves distinct branches of the UPS, distinguished by their dependency on HEI10. This result is consonant with studies showing that DMC1 is the dominant DNA strand-exchange activity during meiosis, while RAD51 plays an essential supporting role that does not require its catalytic activity (*22*, *23*). Differential regulation of DMC1 and RAD51 by the RNF212-HEI10 pathway suggests that RAD51 also plays a later role in meiotic recombination.

We extended our previous studies showing that *Rnf212* and *Hei10* mutations differentially affect turnover of ZMM factor, MutSγ, with faster turnover in the absence of RNF212 and persistence when HEI10 was absent (*20*, *21*)(**fig. S13**). Differential effects of *Rnf212^-/-^* and *Hei10^-/-^* mutations were also observed for MER3 (Fig. 4K,L) and a third ZMM factor, TEX11 (**fig. S13**). In all three cases, persistence of high numbers of ZMM foci seen in *Hei10^-/-^* mutant nuclei was largely RNF212 dependent. We also confirmed that general DSB marker, γH2AX, persists along synapsed chromosomes of *Hei10^-/-^* spermatocytes (*20*); and showed that persistence was dependent on RNF212 (**fig. S13**).

Although crossing over is abolished in *Rnf212^-/-^* and *Hei10^-/-^* mutants, overall DSB repair remains efficient (*20*, *21*). Moreover, unlike pan inhibition of SUMO and ubiquitin conjugation (Fig. 2), *Rnf212^-/-^* and *Hei10^-/-^* mutations do not cause synapsis defects, which could cause secondary defects in recombination. These considerations support direct roles for the SMS and UPS in regulating recombination. Consistently, subsets of RNF212 and HEI10 foci precisely localize to recombination sites (*20*, *21*). However, costaining for general recombination marker, RPA, and either SUMO, ubiquitin or proteasomes revealed only 28-38% colocalization (**fig. S14**). Thus, the bulk of chromosomal SUMO/ubiquitin/proteasomes does not stably localize with recombination sites, suggesting that the branch of the SUMO-UPS pathway defined here may regulate recombination indirectly via chromosome axes. That said, most RPA foci localized immediately adjacent to, or interdigitated with, SUMO/ubiquitin/proteasome signals (fig. S14). Moreover, modification of recombination factors by SUMO/ubiquitin and interaction with proteasomes are expected to be transient.

In conclusion, the SMS and UPS function coordinately to facilitate major transitions of meiotic prophase including axis morphogenesis, homolog synapsis and recombination (**fig. S15**). RNF212 and HEI10 define an axis-associated, SUMO-ubiquitin-proteasome relay that mediates turnover of the subset of recombination factors that act after DMC1-promoted homolog pairing. This stage marks a key regulatory transition during which a small number of crossover sites are designated from the large pool of ongoing recombination events in such a way that each chromosome pair acquires at least one chiasma (**fig. S15**)(*24*). We suggest RNF212-mediated SUMOylation establishes a precondition for this crossover/non-crossover differentiation process by rendering the turnover of key recombination factors contingent on HEI10-dependent proteolysis. At most recombination sites, HEI10-targeted proteolysis predominates to destabilize nascent intermediates and promote a non-crossover outcome. Designation of a crossover outcome ensues at a minority of sites where RNF212-dependent stabilization predominates, enabling crossover-specific events including formation of double-Holliday junctions and recruitment of factors such as MutLγ (*21*). Absent the RNF212-HEI10 pathway, recombination factors that would normally experience transitory stabilization instead turnover more rapidly, crossover designation does not occur and all DSBs are repaired with a non-crossover outcome.

Our model is consonant with the possibility that HEI10 is a SUMO-targeted ubiquitin ligase (STUbL). Indeed, the RNF212-HEI10 pathway shares general features with STUbL pathways that regulate DSB repair in somatic cells (*25*). These pathways stall DSB repair at an early step until appropriate features have been installed that minimize aberrant repair (*26*). We suggest that SUMO conjugation may be a common checkpoint-like mechanism to stall biological processes to enable implementation of important regulatory decisions.

## Acknowledgments

We thank Aldrin Gomes, Bernard de Massy, Christer Höög and Vishva Dixit for antibodies; Michael Paddy for assistance with SIM and the Hunter Lab for support and discussions. This work was supported in part by NIGMS grant GM084955. N.H. is an Investigator of the Howard Hughes Medical Institute.

## Supplementary Materials

Materials and Methods

Figs. S1 to S15

